# The m^6^A Landscapes on the RNA of Mumps Virus and the Host Modulate the Viral Replication and Antiviral Innate Immunity

**DOI:** 10.64898/2026.03.09.710556

**Authors:** Shixuan Wang, Yifei Zhang, Tongyu Ping, Xueping Zeng, Mei Yang, Wanqing Wang, Qing Hu, Ting Fu, Yueqing Chen, Mijia Lu

**Affiliations:** School of Laboratory Medicine and Bioengineering, Hangzhou Medical College, Hangzhou Zhejiang, 310013, China; State Key Laboratory for Conservation and Utilization of Bio-resource and School of Life Sciences, Yunnan University, Kunming Yunnan, 650091, China

**Keywords:** Mumps virus, N6-methyladenosine, viral replication, anti-viral innate immunity

## Abstract

N6-methyladenosine (m^6^A) is the most prevalent internal modification in eukaryotic mRNA and has emerged as a critical regulator of RNA virus infection. However, its role in the mumps virus (MuV), a non-segmented negative-strand (NNS) RNA virus, remains undefined. Here, we comprehensively characterize the m^6^A epitranscriptome of MuV JL2 strain and its functional impact on viral replication and host innate immunity. Using single-base resolution GLORI-seq, we identified abundant m^6^A modifications on MuV genomic, antigenomic and messenger RNAs, with uneven distribution and non-canonical motif enrichment. Genetic depletion of METTL3 enhanced viral replication by facilitating RNA synthesis and nucleocapsid encapsidation in Vero-E6 cells. In epithelial A549 and monocytoid THP1-iDC cells, MuV infection triggered robust type I/III interferon and proinflammatory responses by activating multiple pattern recognition receptors in a m^6^A-dependent manner and induced incomplete iDC maturation characterized by downregulated HLA class II; however, the latter molecule can be partially restored by METTL3-knockdown. The MuV infection reshaped the host m^6^A landscape with a positive correlation between m^6^A enrichment and transcripts enhancement of numerous innate immune genes in a cell type–specific manner. Functional analyses revealed that host m^6^A machinery modulates antiviral signaling, which varies between viral infection and viral RNA-transfection due to the infection-induced host m^6^A machinery fluctuation. Collectively, our findings demonstrate that m^6^A serves as a bidirectional regulator during MuV infection, simultaneously constraining viral replication and modulating host immune recognition, thereby highlighting RNA methylation as a pivotal determinant of MuV pathogenesis and a potential target for optimized immunogenicity.

**Importance:** N6-methyladenosine (m^6^A) modification has emerged as a key regulator of RNA virus infection, yet its role in mumps virus (MuV) remains undefined. Here we map the single-base resolution m^6^A landscape of a recombinant MuV and demonstrate that viral m^6^A restricts genome encapsidation and replication while attenuating innate immune sensing. MuV infection extensively remodels host m^6^A profiles in epithelial and dendritic cells, preferentially enhancing methylation of innate immune transcripts. Strikingly, m^6^A-deficient MuV RNA exhibits increased affinity for RIG-I–like and Toll-like receptors, eliciting stronger interferon and proinflammatory responses. Our findings identify m^6^A as a bidirectional regulator of MuV replication and immunity and suggest an epitranscriptomic strategy to optimize live-attenuated vaccine design.

## Introduction

Mumps virus (MuV), a member of the *Paramyxoviridae* family, is an enveloped, non-segmented, negative-sense single-stranded RNA virus. Its genome is approximately 15.4 kb in length and encodes seven structural proteins: the nucleocapsid protein (N), phosphoprotein (P), matrix protein (M), fusion protein (F), hemagglutinin-neuraminidase (HN), small hydrophobic protein (SH), and the large protein (L). The N protein encapsidates the viral RNA to form a helical nucleocapsid, which is then assembled into the viral envelope. The envelope protein HN and F are the primary targets for neutralizing antibodies^[1]^. The HN mediates viral attachment through recognition of sialic acid receptors and exhibits neuraminidase activity, whereas the F protein facilitates membrane fusion between the viral envelope and the host cells^[2]^. Some attenuated strains of MuV are main component of the measles, mumps, and rubella (MMR) vaccine, which has been implemented in more than 122 countries and has significantly reduced global morbidity and mortality associated with these diseases in the last few decades^[3,4]^. Benefit from the reverse genetics system for MuV, it’s possible to insert other pathogens’ antigen gene into MuV’e genome and thus a quadrivalent vaccine based on MMR can be dosed in one shot. However, the traditional MMR vaccines are administered via intramuscular injection (*im.*) by single-use syringes. In impoverished countries and regions, the shortage of single-use syringes hinders the immunization coverage and brings additional risks such as the epidemic of infectious diseases caused by sharing syringes. During the COVID-19 pandemic, we developed a panel of Measles virus-, MuV- and vesicular stomatitis virus (VSV)-vectored SARS-CoV-2 vaccine candidates, whose convenience and effectiveness of intranasal (*in.*) immunization were validated in animal models, especially in the induction of high-level mucosal immunity, an outcome unachievable with subcutaneous injection^[5,6]^. Also, due to the tropism of respiratory viruses for respiratory epithelial cells, *in.* immunization offers significant advantages and aligns better with the “needle-free” trend. Thus it is necessary to clarify the MuV-induced innate immunity and potential inflammatory responses in lung.

To minimize the virulence and optimize the immunogenicity of a negative-RNA virus, multiple mutation strategies has been applied to inhibit virus RNA-dependent RNA polymerase (RdRP) activity^[7–10]^, and recently we discovered that the N6-methyladenosine (m^6^A) modification in viral RNA is also a potential target for more intense innate and adaptive immune responses in the case of human metapneumovirus (hMPV) and respiratory syncytial virus (RSV)^[11,12]^, whose m^6^A-deficient viral RNA has higher affinity to Retinoic acid-inducible gene I (RIG-I), one of the cytosolic pattern recognition receptors (PRRs) and thus activate more robust type I interferon (IFN) response. m^6^A is one of the most prevalent internal RNA modifications among over 170 known post-transcriptional modifications (PTMs) in eukaryotic cells and viruses. It regulates coding and non-coding RNAs’ nuclear-export, stability, translation, and binding affinity, and helps viral RNA mimic host RNA to evade innate immune surveillance by cytoplasmic pattern recognition receptors (PRRs)^[13]^. However, to date it remains unclear whether there’s m^6^A on MuV RNA and whether this PTM functions on MuV’s propagation or immunogenicity.

In this study, we identified prominent m^6^A peaks on the genomic RNA of MuV JL2 strain using a single-base-resolution m^6^A mapping technique GLORI^[14]^, and explored the epitranscriptomic interplay between MuV and its host cells in regard to the virus replication and innate immune evasion.

## Materials and methods

### Biosafety statement

All experiments with infectious reagents including live virus were conducted under biosafety level 2 (BSL2) at Hangzhou Medical College using standard operating procedures and were approved by the BSL2 Advisory Group and the Institutional Biosafety Committee (IBC).

### Cell lines

A549, Vero (African green monkey, ATCC no. CCL81), Vero-E6 (ATCC CRL-1586) and HEK293T cell lines were purchased from Pricella Biotechnology Co., Ltd. Wuhan, China, and were cultured in Dulbecco’s modified Eagle’s medium (DMEM, Gibco, C11995500BT) supplemented with 10% fetal bovine serum (FBS, Sigma, B7446) and Pen. Strep. (100 IU/mL each) in 37°C incubator with 5% CO_2_. The human leukemia cell line THP-1 is a kind gift from Dr. Yaowei Huang, cultured in RPMI 1640 (Gibco, C11875500BT) supplemented with 10% FBS and Pen. Strep. and used for monocytoid immature dendritic cells (iDCs)-induction according to published methods^[15]^ with modification. Briefly THP-1 cells were seeded in a flask (10^6^ cells /mL) with complete 1640 culture medium, supplemented with recombinant human granulocyte-macrophage colony-stimulating factor (rhGM-CSF) and human interleukin 4 (rhIL-4) (200 ng/mL for each, Novoprptein, C003 and C050, respectively). The culture medium containing the cytokines was refreshed every other day until 6-day post seeding. At day 6 the cells were scraped off the flask and quantified for the following experiments. To obtain FTO-inactivated THP1-iDC, the chemical inhibitor FB23-2 (MCE, HY-127103) was added into the medium (10 μM) at day 4 of the induction for a 2-day treatment.

### METTL3-deficient cells

Three methods were used in our study to downregulate or inhibit METTL3 in multiple cell lines. The first is to infect Vero-E6 cells with shMETTL3 (sense: 5’-TAATTCGTCTGAAGTGCAGC-3’)-expressing lentivirus (Tsingke Biotech Co., Ltd. Beijing China) and screen the infected cells with 5 μg/mL puromycin (Solarbio Life Sciences, P8230) and monoclonal-selection. The second way is to transfect cells with METTL3-targetting siRNA (siMETL3: 5’-CTGCAAGTATGTTCACTATGA-3’, Ribo Life Science Co., Ltd, Suzhou China). The decreased METTL3 expression was then tested by immunoblotting. The third method to inhibit the methyltransferase is to treat cells with STM2457 (MCE, HY-134836), a chemical drug which inhibits METTL3 activity efficiently^[16]^, at 10 μM for 24 h and then evaluate the m^6^A deduction with purified total RNA or mRNA by dot blot.

### Recombinant MuV (rMuV) stock and virion RNA (vRNA) preparation

The MuV strain JL2 used in this study is a recombinant virus (rMuV) rescued with full-length genome plasmid pYES2-MuV-JL2 and assistant plasmids (pN, pP and pL) in our lab as described in our previous work^[6]^. To acquire pure and high-titer virus stock for the following infection and vRNA-extraction, we use the purification method as described^[11]^. Briefly, the culture medium of rMuV-infected Vero cells, containing progeny virus, was harvested, clarified to remove cell debris by centrifugation at 5,000 rpm for 10 min at 4°C. The supernatant was ultracentrifuged with 25% (w/v) sucrose cushion at 25,000 rpm at 4°C for 2 h. The virus pellet was then resuspended with NTE buffer containing 10% (w/v) trehalose (Macklin, D807342), aliquoted and titrated by plaque assay in Vero cells. To acquire m^6^A-defficient rMuV stock, STM2457-treated Vero and Vero-E6-shMETTL3 cells were used to amplify the virus, which was then harvested and purified by ultracentrifugation as described above. The purified virus particle was then lysed by TRIzol reagent (Invitrogen, 15596026CN) for virion RNA (vRNA) extraction according to the product manual.

### m^6^A methylated RNA immunoprecipitation sequencing (m^6^A MeRIP-Seq)

The m^6^A MeRIP-Seq service was provided by CloudSeq Inc. (Shanghai, China). The total RNA of rMuV-infected (MOI= 0.2) or mock-infected THP1-iDC (n=2) was extracted at 16 h post-infection (hpi) and subjected to immunoprecipitation using the GenSeq® m6A MeRIP Kit (GenSeq Inc.), following the manufacturer’s instructions. Briefly, RNA was randomly fragmented to ∼200 nt by RNA Fragmentation Reagents.

Protein A/G beads were coupled to the m^6^A antibody by rotating at room temperature for 1 h. The RNA fragments were incubated with the bead-linked antibodies and rotated at 4°C for 4 h. After incubation, the RNA/antibody complexes were washed several times, and then, the captured RNA was eluted from the complexes and purified. RNA libraries for IP and input samples were then constructed with GenSeq® Low Input Whole RNA Library Prep Kit (GenSeq, Inc.) by following the manufacturer’s instructions. Libraries were qualified using Agilent 2100 bioanalyzer (Agilent) and then sequenced.

### Single-base-resolution m^6^A mapping by GLORI (m^6^A GLORI-Seq)

GLORI sequencing service was provided by Cloud-Seq Biotech (Shanghai, China). The total RNA of rMuV-infected (MOI= 0.2) or mock-infected A549 (n=2) was extracted at 20 hpi and treated with GenSeq® GLORI kit (GenSeq Biotech Inc., Shanghai) according to the manufacturer’s instruction. Briefly, RNA was incubated with fragmentation buffer for 4 min at 94°C, then glyoxal solution and DMSO were added, and the mixture was incubated for 30 min at 50℃ in a preheated thermocycler. The samples were cooled down to room temperature, mixed with saturated H3BO3 solution, and incubated at 50°C for 30 min, followed by the addition of deamination buffer and 8 h incubation at 16°C. Finally, the RNA was purified by ethanol precipitation, and sequencing libraries were constructed with GenSeq® eCLIP RNA Library Prep Kit (GenSeq Biotech Inc., Shanghai) following the manufacturer’s instructions. The constructed sequencing libraries were subjected to quality control and quantification by the BioAnalyzer 2100 system (Agilent Technologies, USA), followed by 150 bp double-end sequencing.

### Whole transcriptome sequencing

The whole transcriptome of the infected and mock-infected THP1-iDC was performed by CloudSeq Inc. using the Input samples for m^6^A MeRIP-Seq (n=2). RNA sequencing service was provided by CloudSeq Inc. (Shanghai, China). Briefly, ribosomal RNA (rRNA) was removed from samples using GenSeq® rRNA Removal Kit (GenSeq, Inc.), and the rRNAdepleted samples were used for library construction with GenSeq® Directional RNA Library Prep Kit (GenSeq, Inc.) by following the manufacturer’s recommendations. The RNA was fragmented into ∼300 nt in length. The first-strand cDNA was synthesized from the RNA fragments by reverse transcriptase and random hexamer primers, and the second-strand cDNA was synthesized in the 2^nd^ Strand Synthesis Buffer with dUTP Mix. Then, the double-stranded cDNA fragments were subjected to end-repair and dA-tailing, followed by adapter ligation. The adapter-ligated DNA samples were PCR-amplified and purified to generate sequencing libraries. Finally, the libraries were sequenced with a sequencer on the paired-end 150 bp mode.

### mRNA sequencing

The total RNA sample (A549, n=3), including the diplicates for m^6^A GLORI-seq and a third samples prepared together with the other two at the same time with same way, were used for transcriptome (mRNA) sequencing by Majorbio Bio-pharm Biotechnology Co., Ltd. (Shanghai, China). RNA purification, reverse transcription, library construction and sequencing were performed according to the manufacturer’s instructions (Illumina, San Diego, CA). The RNA-seq transcriptome library was prepared following Illumina® Stranded mRNA Prep Ligation from Illumina (San Diego, CA) using 1μg of total RNA. Briefly, messenger RNA was isolated according to the polyA selection method by oligo(dT) beads and then fragmented by the fragmentation buffer firstly. Secondly double-stranded cDNA was synthesized using a SuperScript double-stranded cDNA synthesis kit (Invitrogen, CA) with random hexamer primers (Illumina). Then the synthesized cDNA was subjected to end-repair, phosphorylation and ‘A’ base addition according to Illumina’s library construction protocol. Libraries were size-selected for cDNA target fragments of 300 bp on 2% Low Range Ultra Agarose followed by PCR amplification using Phusion DNA polymerase (NEB) for 15 PCR cycles. After quantification by Qubit 4.0, the paired-end RNA-seq sequencing library was sequenced with the NovaSeq X plus sequencer (2 × 150 bp read length).

### RNA quantification by reverse transcription-real time PCR (RT-qPCR)

To quantify the copy number of the viral genomic RNA (gRNA, negative strand) and antigenomic RNA (agRNA, plus strand) in vRNA, leader-primer and trailer-primer, respectively, were used to generate a long cDNA strand in One-step cDNA synthesis mix (TransGen Biotech, AU311), followed by qPCR (Vazyme, Q311-03) with *HN* gene-specific primers. The 10-fold serial dilutions of pYES2-MuV-JL2 (10∼10^9^ copies/μL) were used for standard curve generation and RNA copies calculation. To quantify the transcript mRNA of host cells and virus, the total RNA was extracted from cells with TRIzol reagent at the indicated hpi or hours post-transfection (hpt). The cDNA pool was generated with random primer (for host transcripts) or anchored oligo (T)_23_ (for viral mRNA, customer-synthesized by Tsingke Biotech), and qPCR was performed with the indicated primer pairs with internal control of *β-actin*. All the primers used in this study are listed in Table 1.

**Table 1.**
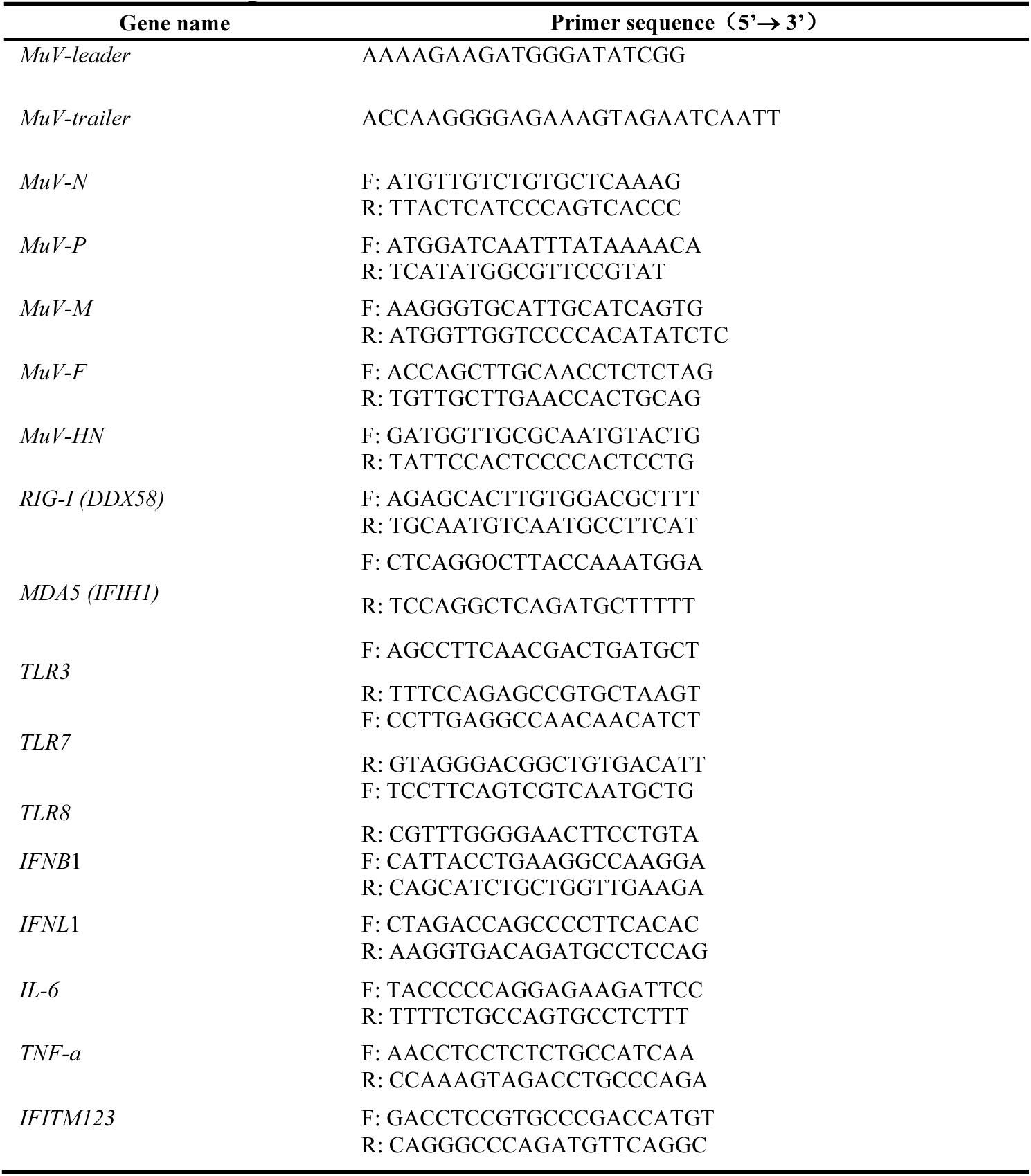
Primer sequences.

### Viral encapsidation evaluation by RNase digestion

Vero-E6-shMETTL3 or Vero-E6-shN cells were infected with rMuV at an MOI of 0.5. At 24 and 48 HPI the infected cells were lysed by Pierce IP lysis buffer (ThermoFisher, 87787) on ice for 20 min. The cell lysate containing the propagating virus nucleocapsid was clarified by 12,000 rpm for 10 min at 4°C and aliquoted for RNase-treatment. In a 50 μL reaction system containing 10 mM Tris-HCl (pH 7.4), 15 mM NaCl, 1 mM EDTA, 30 μL cell lysate aliquots were treated with the combination of 5 μg RNase A (Beyotime, ST576) and 1 U RNase III (Beyotime, R7086), with the control of PBS, incubated at 37°C for 30 min. The RNase was then inactivated by 50 mM EDTA and the reaction system was immediately subjected to RNA extraction by the viral RNA-extraction kit (Transgen, ER211). All the extracted RNA was quantified for gRNA and agRNA by RT-qPCR as described above. The RNA copies post RNase-treatment is taken as encapsidated by N protein, and the ratio of RNase-/PBS-treatment according to the Ct value was calculated as the relative encapsidated viral RNA.

### RNA dot blot

To test the m^6^A level the vRNA was denatured at 98°C for 10 min, immediately cooled down on ice, mixed with 20 × SSC buffer and dropped on the Amersham Hybond-N+ membrane (Cytiva, RPN203B) with the indicated amount. The RNA sample was allowed to air dry followed by UV-link for twice, and then subjected to BSA blocking, rabbit anti-m^6^A monoclonal antibody (CST, 56593) and HRP-conjugated Goat anti-Rabbit IgG (Proteintech, SA00001-2) incubation, and development in the substrate (ThermoFisher, 34580) for dots visualization. The membrane was stained with methylene blue solution (0.1%, w/v) for loading amount control.

### RNA-Transfection

For siRNA-introduced gene-knockdown, the cells in 24-well plate were transfected with gene-specific siRNA (30 pmol/well), controlled with the same amount of non-specific siRNA (siN), with the assistance of Lipofectamine RNAiMAX (ThermoFisher, 13778100) 3-day prior to the subsequent handling. To introduce vRNA into the target cells, the mixture of vRNA (10^6^ copies/well) and transfection reagent Golden-R (Golden Transfer, 190425015) was added to the culture medium of A549 (2×10^5^ cells/ well) or THP1-iDC (4×10^5^ cells/ well) in 24-well plate, controlled with the same volume of phosphate buffered saline (PBS). The transfected cells were subjected to the following infection or collection at the indicated time points for further determination by WB, RT-qPCR, and flow cytometry.

### rMuV titration by plaque assay and immune staining

Confluent Vero cells in 24-well plates were infected with 10-fold serial dilutions of rMuV (125 µL/well) by inoculation at 37°C for 1.5 h with constant gentle shaking. The cells were then covered with 0.5 mL DMEM overlay containing 0.8% (w/v) methylcellulose, 0.12% (v/v) NaHCO_3_, 1% (w/v) FBS, 25mM HEPES, 2mM L-Glutamine, 100 µg/mL streptomycin and 100 U/mL penicillin, and cultured at 37°C. At 3-day post inoculation (dpi) cells were fixed with 10% (v/v) neutral-buffered formalin (NBF) at room temperature for 30 min. The overlay was then removed together with NBF, and the rMuV plaques were visualized by immune staining described previously^[11]^. Briefly, the fixed cells were washed with PBS and permeabilized in 0.4% (v/v) Triton X-100 at room temperature for 10 min, blocked with 1% (w/v) bovine serum albumin (BSA, Macklin, B824162) for 1 h, incubated successively with anti-MuV N protein monoclonal antibody (abcam, ab9880) at 4°C overnight and horseradish peroxidase (HRP)-labelled goat anti-mouse secondary antibody (SinoBiological, SSA007) at 37°C for 1h, then incubated with chromogen substrate of AEC (3-amino-9-ethylcarbazole, Merck, A5754). The rMuV N protein-expressing cell clusters were developed as red-brown dots and countered under a microscope for viral titer (plaque-forming units per mL, PFU/mL) calculation.

### Immunoblotting (Western blot, WB)

Cells were lysed in RIPA buffer (Beyotime Biotechnology, P0013K) supplemented with protease inhibitor cocktail (Sigma-Aldrich, 11873580001) on ice followed by 12,000 rpm centrifugation at 4°C and denatured with 5× loading dye at 98°C for 10 min. The protein samples were isolated by SDS-PAGE and transferred onto a nitrocellulose (NC) membrane (Cytiva, Cat. No. 10600002). The protein-bound membrane was then blocked with 5% (w/v) skim milk, incubated with primary antibody at 4°C overnight followed by horse radish peroxidase (HRP)-conjugated secondary antibody, and then developed with the substrate (ThermoFisher, 34580). The primary and secondary antibodies used in this study are listed in Table 2.

**Table 2.** Antibodies in this study.

| Antibody | Supplier, Cat. No. |
| --- | --- |
| Mouse anti-Mumps nucleoprotein antibody | Abcam, ab9880 |
| Mouse anti-MAVS monoclonal antibody | Proteintech, 66911-1-Ig |
| Rabbit anti-RIG-I recombinant monoclonal antibody | Abcam, ab180675 |
| Rabbit anti-β-actin recombinant monoclonal antibody | Proteintech, 81115-1-RR |
| Rabbit anti-FTO polyclonal antibody | Proteintech, 27226-1-AP |
| Rabbit anti-ALKBH5 polyclonal antibody | Proteintech, 16837-1-AP |
| Rabbit anti-TLR3 monoclonal antibody | Cell Signaling, 6961 |
| Mouse anti -METTL3 monoclonal antibody | Proteintech, 67733-1-Ig |
| Rabbit anti -METT14 polyclonal antibody | Proteintech, 26158-1-AP |
| Rabbit anti-pIRF3(S386) monoclonal antibody | Abcam, ab76493 |
| Rabbit anti-pIRF7(S477) monoclonal antibody | Cell Signaling, 12390 |
| Rabbit anti-β-actin recombinant monoclonal antibody | Proteintech, 81115-1-RR |
| Goat anti-Rabbit IgG (H+L) (HRP) | Proteintech, SA00001-2 |
| Goat anti-mouse IgG (HRP) | SinoBiological, SSA007 |

### RNA immunoprecipitation (RIP) assay for the affinity of PRR molecules to virion RNA

HEK293T cells cultured in 6 cm dish with Pen. Strep.-free DMEM were transfected with the mixture of 10 μL Lipofectamine 3000 and 5 μg Flag-tagged RIG-I-, MDA5-, TLR3- or TLR7-expressing plasmid with the control of vector plasmid, and then cultured for another 24 h. The cells were then transfected with MuV virion RNA (100 copies/ cell) and were lysed with Pierce lysis buffer (Thermofisher, 87787) supplemented with RNase inhibitor (NEB, M0314, 0.2 U/μL lysate) at 2 hpt. One tenth of the cell lysate supernatant (Input) was used for overexpression confirmation by immunoblotting, 3/10 of the Input was used for vRNA Input, and the rest was incubated with Pierce™ anti-DYKDDDDK (Flag) magnetic agarose (ThermoFisher, A36797) at room temperature for 30 min. The agarose beads were washed and the bound protein was released from the beads by another 10 min incubation with 3× Flag peptide (1.5 mg/mL). The resulting 1/10 I.P. elution was used for target protein detection by WB and the rest, together with the RNA Input, were subjected to RNA extraction for vRNA quantification by RT-qPCR.

### Flowcytometry

MuV-infected or vRNA-transfected THP1-iDCs were harvested and washed with PBS by centrifugation at 250 ×g for 5 min, and stained with Horizon™ Fixable Viability Stain 510 (FVS510, BD Pharminge, 564406) for dead cells exclusion, and then stained with fluorescent-labeled antibodies targeting DCs’ surface marker CD11c, CD80, CD83, CD86, CD274, HLA-ABC and HLA-DR,DP,DQ (BD, 56256, 751734, 558017, 740990, 551073, 740407, 550853, respectively) in staining buffer (PBS containing 1% FBS). The stained cell suspension was then subjected to flow cytometry, the fluorescence strength of each single surface marker was generated by the software FlowJo (BD, v.10.8.1) and calculated as mean ± standard deviation (SD).

### Statistical analysis

Statistical analysis was performed by Student’s t test, one-way or two-way ANOVA and multiple comparisons are indicated in the figure legend. p values of less than 0.05 were considered statistically significant, *:*p* < 0.05, **:*p* < 0.01, ***:*p* < 0.001, ****:*p* < 0.0001.

## Results

### MuV RNA carries N6-methyladenosine (m^6^A) modification

To date m^6^A modifications have been identified in the RNA of numerous viruses, such as SARS-CoV-2, Zika virus (ZIKV), West Nile virus (WNV), dengue virus (DENV), hepatitis C virus (HCV), simian virus 40 (SV40), and HIV^[17–20]^, as well as multiple non-segmented negative strand (NNS)-RNA viruses in our previous work^[11,21,22]^. To reveal whether MuV ‘s RNA also carries m^6^A and whether the infection impacts the m^6^A landscape of the host’s transcriptome, the total RNA of rMuV-infected A549 cells were collected for GLORI m^6^A sequencing. Benefited by the single-base resolution of the sequencing technique, there’re 178 and 328 confident m^6^A peaks (m^6^A ratio > 0.1) identified on the negative (-) strand (viral gRNA) and the plus (+) strand (agRNA and mRNA), respectively (Fig.1a, *Supplementary* data Excel file 1). In each gene and intergenic region m^6^A peaks were detected in single-base resolution with various depths and coverage ratio, and the gray column of A and T indicates a 100% m^6^A modification in the sense and antisense strand, respectively (Fig. 1a, panel 1∼4). The highest ranked m^6^A motif patterns in viral RNA were concluded as MAACA and UAAAGW (M= A/C, W= A/U), which don’t match the canonical eukaryotic m^6^A motifs RRACH (R= A/G, H= A/C/U) (Fig.1b). Among the hundreds of m^6^A peaks there’re 45 and 83 peaks presented in both samples’ (-) and (+) strand, respectively. From these relatively stable peaks located in both samples, 14 on (-) strand and 30 on (+) strands with high confidence (m^6^A ratio > 0.2) were listed in Table 3. On the (-) strand most of the high-confidence peaks are located in HN (5/14) and N (3/14) genes, while on the (+) strands the peaks were most enriched in genes N and P (7/30 for each), followed by F gene (6/30), reflecting an unequal m^6^A distribution character which was also found in human, yeast, and other viruses transcripts^[18,23–25]^. Importantly rMuV’s propagation was upregulated in METTL3-knockdown (shMETTL3) Vero-E6 cells (Fig.1c) according to plaque assay-titrated infectious viral particle production and RT-qPCR-measured viral gRNA and agRNA (Fig.1d, e). Consistently the viral mRNA copies of *n*, *p*, *f*, and *m* genes were also enhanced in the shMETTL3-Vero-E6 (Fig.1f). As the infected Vero-E6 cell lysate was treated with RNases, the nascent and unencapsidated gRNA is supposed to be cleaved^[26]^, the uncleaved vRNA thus is calculated as encapsidated by N protein, and the ratio of RNase-treated / PBS-control is an indicator for encapsidation efficiency. Interestingly we find that not only gRNA but also agRNA was reserved post RNase treatment according to the relative encapsidation of 0.1∼0.2, which is less than that of gRNA (0.2∼0.4), except for that in the METTL3-depleted cells at 24 hpi (*p*> 0.05, Fig.1g, left panel), suggesting that agRNA is not completely excluded from the nucleocapsid assembly of MuV. Moreover, the encapsidation of both RNA strands was promoted in METTL3-deficient cells (Fig.1g, right panels), implying a suppressive function of m^6^A during MuV nucleocapsid assembly. Thus we for the first time demonstrated that MuV’s rival RNA is m^6^A-modified, which negatively modulates the virual replication in an innate immunity-deficient host cells.

**Fig. 1.**
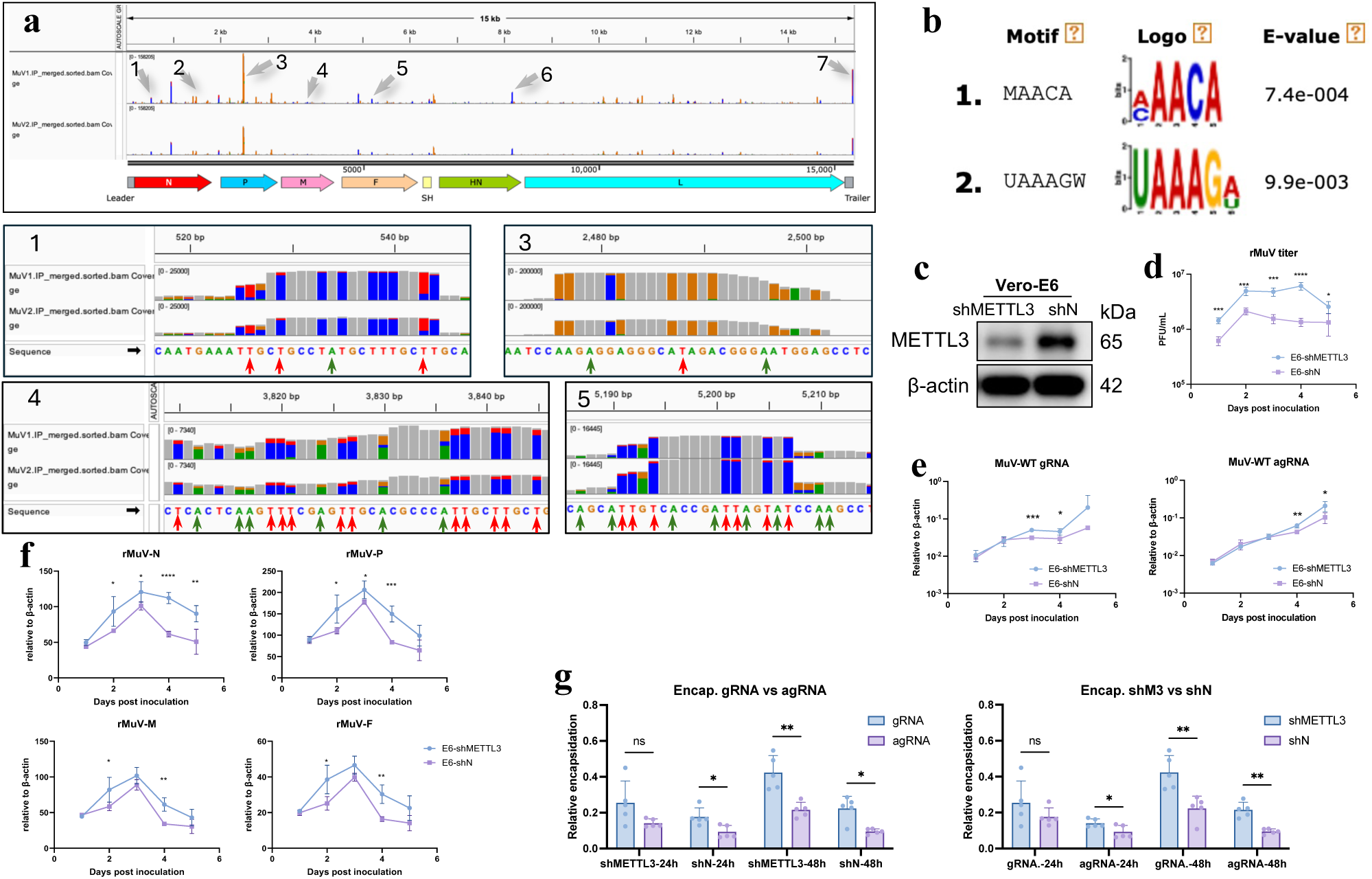
The viral RNA of recombinant MuV JL2 is modified with m^6^A, which hinders the viral replication. **a.** The total RNA of rMuV (JL2 strain)-infected A549 cells was subjected to the high-through m^6^A-sequencing GLORI (n=2), and the resulted m^6^A landscape is visualized by software IGV in the direction of antigenomic RNA (agRNA, plus strand). The presentative m^6^A peaks in dividual genes are labeled with gray arrows (top panel) and zoomed into single-base resolution (middle and bottom panels. Red “T” under red-blue column : partially m^6^A-modified bases in negative strand-gRNA. Green “A” under green-yellow column: partially m^6^A-modified bases in plus strand-agRNA. Green “A” under gray column: m^6^A-modified bases with 100% confidence in agRNA. The green and red arrows indicate the presentative but not all m^6^A sites). **b**. The m^6^A motifs concluded by GLORI-seq with E-values (E-values = the enrichment p-value ✕ the number of candidate motifs tested. The enrichment p-value is calculated using Fisher‘s Exact Test for enrichment of the motif in the positive sequences). **c-f.** The Vero-E6 cells was used to construct METTL3-knockdown cells by shMETTL3-lentivirus infection and puromycin screen. The shMETTL3 cells were tested for decreased METTL3 by immunoblotting, with the control of shN cells (**c**), and then were infected by rMuV at the MOI of 0.2 for dynamic monitoring of alive virus titer (**d**), viral RNAs including gRNA, agRNA (**e**), and mRNAs of multiple genes (**f**). **g.** Vero-E6-shMETTL3 or –shN cells were infected with rMuV at the MOI of 0.5 and collected at 24 and 48 hpi for RNase-treatment, with the control of PBS-treatment. The ratio of RNase-/PBS-treatment according to the Ct value was calculated as the relative encapsidation of viral RNA, which was presented as the comparisons of vRNA strands (left panel) and METTL3 scarcity (right panel). The above data (except for sequencing data) are from independent three experiments and the representative one is presented (**d-g;** mean ± S.D.). ns: *p* > 0.05, \**p* < 0.05, ** *p* < 0.01, *** *p* < 0.001, **** *p* < 0.0001 (Student’s T test, one-way ANOVA, and two-way ANOVA).

**Table 3.**

| Strnad | Sites | MuV1.m<sup>6</sup>A.ratio | MuV2.m<sup>6</sup>A.ratio | Gene name |
| --- | --- | --- | --- | --- |
| gRNA<br>(-) | 525 | 0.31008 | 0.30216 | N |
|  | 526 | 0.7367 | 0.68101 | N |
|  | 543 | 0.6721 | 0.65837 | N |
|  | 1961 | 0.65514 | 0.6214 | N-P |
|  | 4875 | 0.28767 | 0.42857 | F |
|  | 6044 | 0.22368 | 0.2 | F |
|  | 6474 | 0.20588 | 0.23077 | SH-HN |
|  | 7156 | 0.22222 | 0.29412 | HN |
|  | 7160 | 0.26087 | 0.24038 | HN |
|  | 7166 | 0.4574 | 0.47244 | HN |
|  | 7748 | 0.2973 | 0.36364 | HN |
|  | 7749 | 0.42105 | 0.36364 | HN |
|  | 13395 | 0.25 | 0.27273 | L |
|  | 14677 | 0.21111 | 0.24658 | L |
| agRNA<br>and<br>mRNA<br>(+) | 350 | 0.22751 | 0.22619 | N |
|  | 921 | 0.2093 | 0.30435 | N |
|  | 1154 | 0.275 | 0.23006 | N |
|  | 1357 | 0.22222 | 0.22222 | N |
|  | 1528 | 0.24786 | 0.23077 | N |
|  | 1530 | 0.25481 | 0.43966 | N |
|  | 1736 | 0.2766 | 0.36842 | N |
|  | 2144 | 0.60366 | 0.62366 | P |
|  | 2147 | 0.20468 | 0.2 | P |
|  | 2496 | 0.67115 | 0.68421 | P |
|  | 2497 | 0.28932 | 0.28035 | P |
|  | 2759 | 0.28593 | 0.27401 | P |
|  | 2760 | 0.2849 | 0.22162 | P |
|  | 2885 | 0.2139 | 0.25743 | P |
|  | 4157 | 0.23404 | 0.3 | M |
|  | 4255 | 0.2381 | 0.27273 | M |
|  | 4529 | 0.24074 | 0.2 | M-F |
|  | 4643 | 0.35398 | 0.27778 | F |
|  | 4646 | 0.27143 | 0.28846 | F |
|  | 4995 | 0.38158 | 0.26087 | F |
|  | 5792 | 0.20155 | 0.28205 | F |
|  | 6083 | 0.29114 | 0.21053 | F |
|  | 6094 | 0.20833 | 0.35294 | F |
|  | 6221 | 0.41304 | 0.28571 | F-SH |
|  | 7476 | 0.27027 | 0.28846 | HN |
|  | 7479 | 0.52113 | 0.6 | HN |
|  | 11012 | 0.52995 | 0.57692 | L |
|  | 12025 | 0.23636 | 0.28571 | L |
|  | 14630 | 0.41695 | 0.33951 | L |
|  | 15213 | 0.32308 | 0.38776 | L |

### The rMuV induced a significant innate immune response in A549 and THP1-iDCs

MuV is capable of infecting cells in the salivary glands, lungs, brain, and nasal mucosa in macaques, and was proved to reproduce in multiple cell lines including epithelia (Vero, 293G and A549)^[27]^, monocyte (THP-1, U937), T cells (Jurkat and CEM), and B cells (Ramos and Daudi)^[28]^. The high-throughput transcriptome sequencing data revealed that the MuV-infection resulted significant innate immune response in A549 and THP-1-derived iDCs (THP1-iDC) at 16 hpi, according to the enrichment of immune/ defense response- and signaling-related terms in the top 20 upregulated Gene Ontology (GO) items (Fig.2a). The infection activated dramatic type I and type III interferons (IFNs), proinflammatory cytokines and numerous interferon-stimulated genes (ISGs) including IFITs, IFITMs, and multiple PRRs in both cell lines, and enhanced transcription of DC’s maturation biomarkers like CD11c, CD80, and CD86 (Fold change > 2, *p*< 0.05, Fig.2b, *Supplementary* files: A549−rMuV−inn..xlsx, THP1−iDC−rMuV−inn.xlsx). In A549 cells the upregulated typical biomarkers of innate immunity, such as IFN-β, IFN-α, IL-6, TNF-α, and IFITMs, were confirmed by RT-qPCR at multiple time points of 10, 16, and 24 hpi (Fig.2c). The phosphorylated interferon regulatory factor 3 and 7 (pIRF3 and pIRF7), as well as the enhanced RIG-I and MDA5 expression were also detected by immunoblotting (Fig.2d), indicating the MuV-triggered innate immunity and the promoted RLRs expression by the positive feedback of type I IFN secretion. In THP1-iDCs the transcriptomic data revealed innate immune response was similarly confirmed by RT-qPCR and immunoblotting (Fig.2e, f), and moreover the upregulated maturation-related CDs molecules were confirmed by flow cytometry, as well as CD274 (also known as PD-L1), which is induced by type I/II IFNs to limit overproduction of inflammatory cytokines (Fig.2g). Though HLA class I molecules were identified as no change (fold change< 2, or *p*> 0.05) in the sequencing data for 16 hpi (Fig.2b, bottom panel), they were significantly enhanced at 24 and 48 hpi by flow cytometry, in which HLA class II molecules were evaluated as suppressed (Fig.2g). Furthermore the MuV-infection induced deferent RLRs and TLRs response to type I IFN secretion, as in A549 cells only RIG-I, MDA5 and TLR3 were upregulated, but in THP1-iDC all the detected RLRs and TLR3/7/8 transcription was dramatically upregulated (Fig.2h), implying a more predominant function of TLRs in iDCs than in A549. Thus by infection, rMuV is capable of initiating significant innate immunity in epithelial A549 and monocytoid THP1-iDC; however, the downregulated HLA class II molecules indicate the possibility of insufficient DC maturation to impair antigen presentation for CD4^+^ T cell priming, which was also observed in some viral infection cases that resulted in T cell anergy^[29,30]^.

**Fig. 2.**
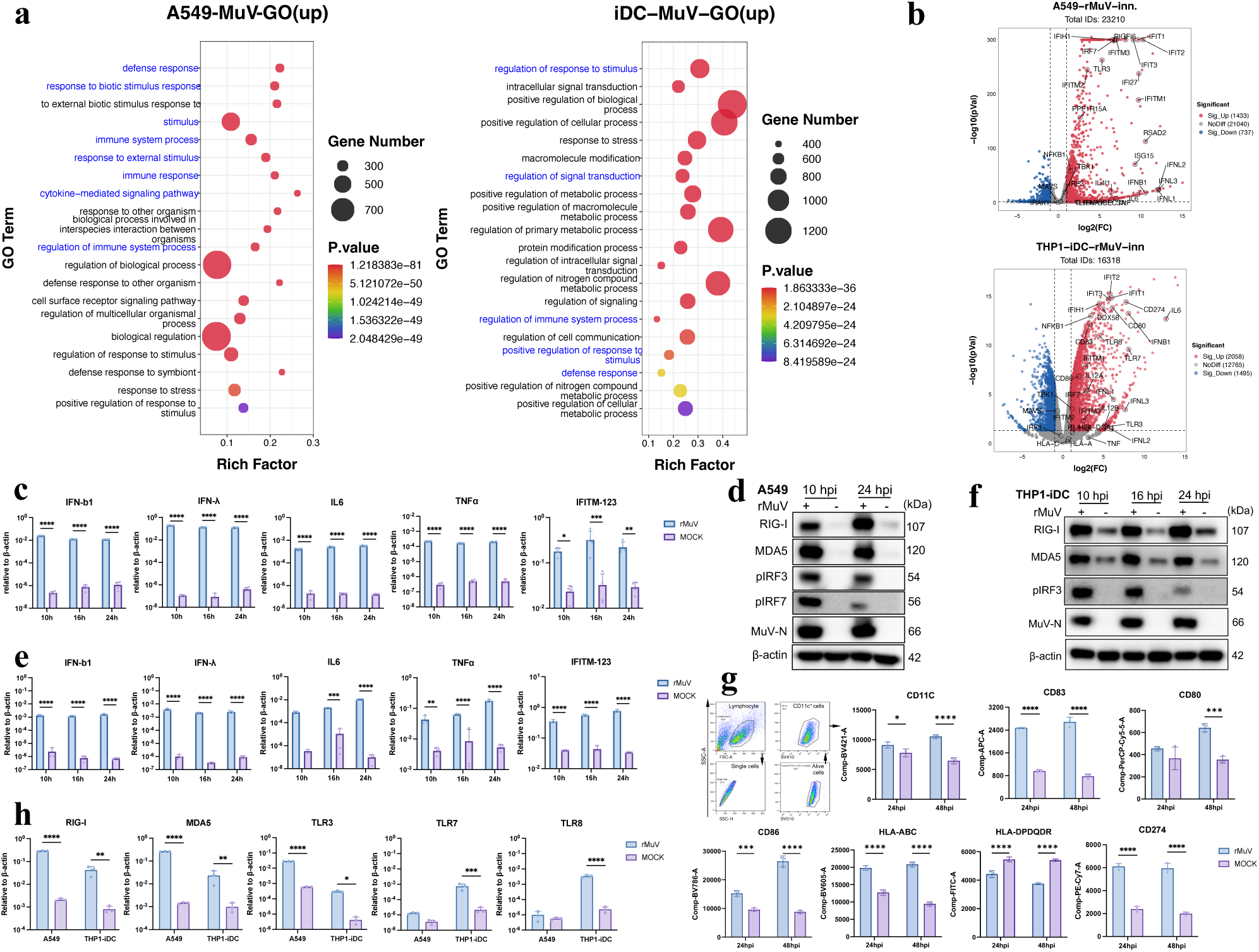
The rMuV induced robust innate immune response in both A549 and THP-iDC cells. **a.** The rMuV-infected (MOI=0.2) A549 (n=3) and THP1-iDC (n=2) were collected at 16 hpi for total RNA extraction and transcriptome sequencing. With the control of mock-infected RNA samples the top 20 upregulated Gene Ontology (GO) items (sorted by *P* values) were listed for each cell type, the innate immune/ defense-response and signaling-related items were marked in blue. **b**. The transcription of innate immunity-related genes (*e.g.* interferons, proinflammatory cytokines, interferon-stimulated genes, and multiple PRRs), and DC’s maturation biomarkers were indicated in the volcano plots (threshold for significance: log2 fold-change > 1, and *p*< 0.05 ). **c-d.** The A549 cells were infected with rMuV at the MOI of 0.2. At 10, 16, and 24 hpi the RNA and cell lysate were collected for transcription determination by RT-qPCR (**c**), and expression of innate immunity-related proteins and viral protein by immunoblotting (**d**). **e-f.** The THP1-iDC were infected with rMuV at the MOI of 0.2, and the innate immunity-related transcripts and proteins were determined by RT-qPCR (**e**) and immunoblotting (**f**), respectively, at 10, 16, and 24 hpi. **g.** The rMuV-infected THP1-iDC (MOI=0.2) were evaluated for maturation by flow cytometry at 24 and 48 hpi. The cells were gated for lymphocyte-single cells-alive cells-CD11c-positive cells and then sub-gated for the indicated molecule-positive cells portions. The fluorescence strength for each single surface marker was generated by the software FlowJo (BD, v.10.8.1). **h.** The cytosolic PRRs of rMuV-infected A549 and THP1-iDC (MOI=0.2) were quantified by RT-qPCR at 16 hpi. The above data (except for sequencing data) are from independent three experiments and the representative one is presented (**c, e, g, h;** mean ± S.D.). \**p* < 0.05, ** *p* < 0.01, *** *p* < 0.001, **** *p* < 0.0001 (Two-way ANOVA).

### The rMuV-infection alters host cells’ m^6^A-landscape

Virus doesn’t possess its own m^6^A machinery but grabs its host’s m^6^A resources, leading to a m^6^A machinery fluctuation and host m^6^A landscape alteration, including global quantity and location shift^[31,32]^. The rMuV-infection led to obviously decreased expression of host’s METTL3, METTL14, and FTO since 10 hpi (A549) or 16 hpi (THP1-iDC), and even a dramatic decrease of ALKBH5 in iDC (Fig.3a), implying an impaired host m^6^A machinery. From the GLORI-seq m^6^A data of A549, the most confident m^6^A motif is concluded as GSAGR and is conserved in the MuV-infected cells, but the less confident motifs altered from YGAGG and GGAGY to AGASG, GGAYG, and GGAGC upon the infection (Fig.3f, two left panels). There’re 3567 gained m^6^A sites were discovered in the infected A549, with only 1164 conserved and 2451 lost sites compared with the mock-infected cells (Fig.3c, upper panels), despite of the METTL3 and METTL14 loss. Within numerous genes the m^6^A location shifted upon infection and statistically resulted in percentage reduction in CDS, but increase in start codon and stop codon (Fig.3d, two left panels). In regard to THP1-iDC, MeRIP-seq data revealed a conserved most enriched m^6^A motif GGACW but altered lower-ranked ones due to the rMuV-infection (Fig.3b, two right panels). Besides 5642 gained m^6^A sites in mRNA, there’re 2664 and 569 unique sites detected in long noncoding RNA (lncRNA) and circle RNA (circRNA), respectively (Fig.3c, lower panels), and similar to A549 the gained m^6^A sites in all RNA types are more than the lost ones, which is different with the massive m^6^A loss upon SARS-CoV-2 variants infection^[32]^. Meanwhile, the rMuV-infection resulted in an enhanced m^6^A percentage in CDS and suppressed it in start codon (Fig.3d, two right panels), and no matter what the location is, most of the m^6^A’s fold-change in THP1-iDC is positively correlated (*p*<0.0001) with the transcript’s quantity fluctuation (Fig.3e, *Supplementary Fig.1a*), suggesting the prominent effect of m^6^A on promoting transcription or stabilizing mRNA. The KEGG analysis, performed separately with m^6^A enriched mRNA or input mRNA, revealed 6 shared pathways in top15 of the subclass “Organismal Systems” according to *p*-values sorting, and 5 out of this 6 shared pathways were typically involved in innate immunity such as NF-κB, TNF, Toll-like receptor, NOD-like receptor, and RIG-I like receptors (RLRs) signaling pathway (Fig.3f). A similar correlative pattern between m^6^A enrichment and enhanced mRNA is also found in A549, though these two sets of sequencing data were carried out with same samples by two companies (*Supplementary Fig.1b*). To assess the relationship between m^6^A modification and transcription/ translation of innate immunity-involved genes, the m^6^A-seq data were visualized by the software IGV. It’s interesting to find that upon infection the modification of RIG-I, MDA5 and TLR3 transcripts is dramatically upregulated in both cell lines, but the enhanced m^6^A on TLR7 and TLR8 transcript is only observed in THP1-iDC (Fig.3h). Combining with the transcription of these PRRs (Fig.2h), the infection-induced m^6^A-deposit seems contribute to the upregulated transcripts in a cell type-dependent manner. This asymmetric regulation pattern is also reported as different PRRs predominance in epithelial cells and monocyte-derived cells in regard to innate immunity initiation^[33,34]^. Also the upregulated IFNs, pro-inflammatory cytokines, ISGs, and iDC maturation markers (Fig.2c, e, g, h) in both cell lines are predominantly m^6^A modified in various regions depending on genes, except for TNF-α whose m^6^A was not detectable in A549, and upregulated mildly in THP1-iDC upon infection (*Supplementary Fig.1c, d*). Collectively, using two m^6^A-seq techniques we demonstrated that rMuV infection initiated a cell type-specific deposit and redistribution of m^6^A, which predominantly modulated the fluctuated expression of numerous genes in a manner of positive correlation, including the key genes contributing to innate immunity signaling.

**Fig. 3.**
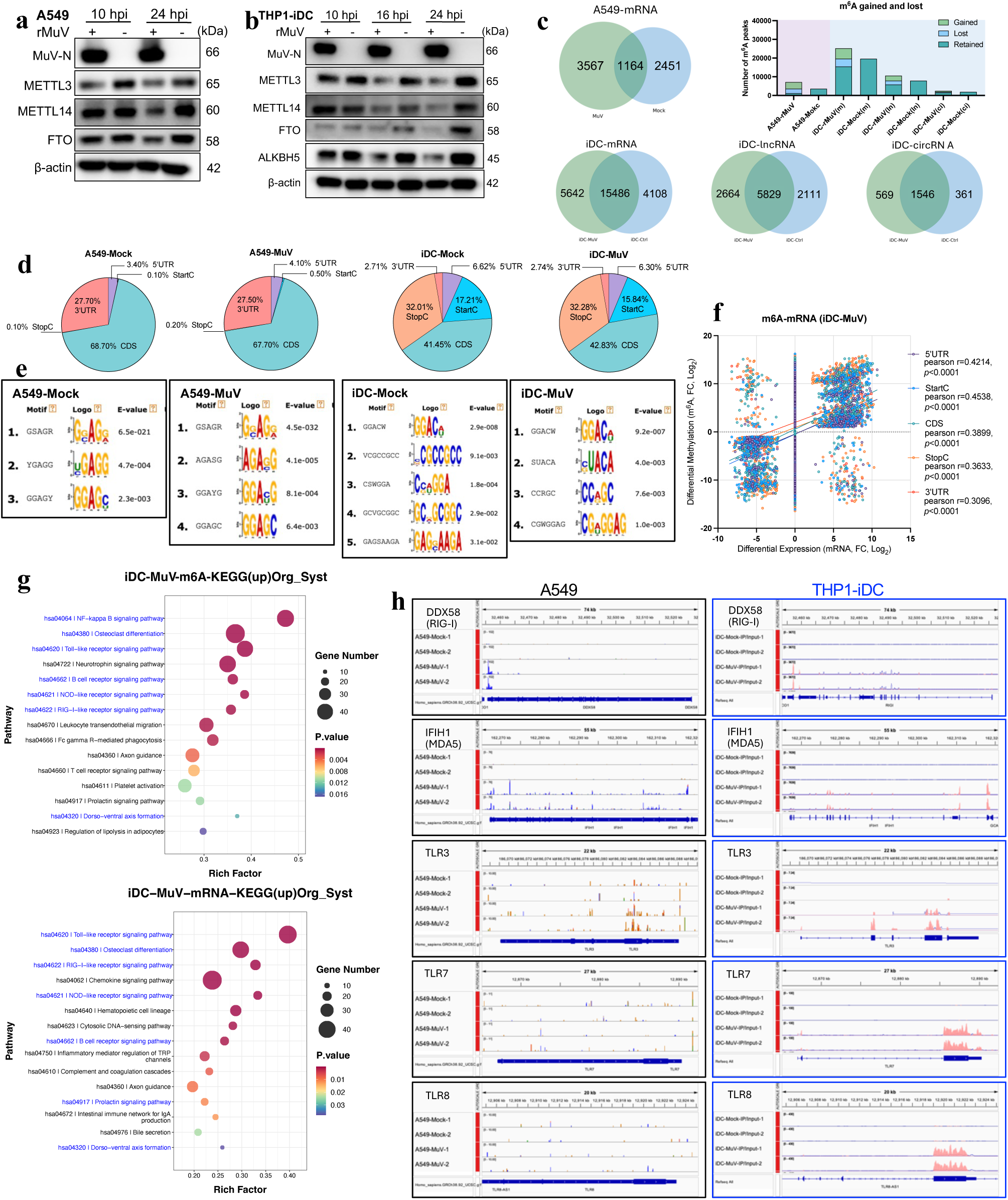
The rMuV-infection reshaped the host m^6^A landscape. **a-b.** A549 (**a**) and THP-iDC (**b**) were infected by the rMuV (MOI=0.2) and the m^6^A machinery proteins were determined by immunoblotting from 10 to 24 hpi. **c-h.** The total RNA of rMuV-infected A549 and THP1-iDC (MOI=0.2) were collected at 16 hpi (n=2) and subjected to m^6^A mapping by GLORI and MeRIP sequencing, respectively. The gained, retained, and lost m^6^A sites in mRNA (A549) and the whole transcriptome (iDC) due to the infection are presented in the Venn plots and a stacked column chart on the upper right (**c**). The distribution frequence of m^6^A site was calculated as percentage (%) for 5’UTR, start codon, CDS, stop codon, and 3’UTR (**d**), the correlation between the m^6^A fold-change (log2) and the mRNA fold-change (log2) in the infected iDC is presented in the scattered plot (**f**). The concluded m^6^A motifs with the top enrichment is listed (**f**). In iDC the most enriched and upregulated KEGG pathways (top 15) in the category “Organismal Systems” are screened from the m^6^A-enriched (top panel) and the Input (bottom panel) mRNA according to the *p* values, the shared pathways in both datasets are labeled in blue (**g**). The m^6^A enrichment in the indicated PRRs transcripts were visualized in the software IGV, with a normalized confidence scale for both infected and mock samples (**h**).

### Host cells’ m^6^A-machinery is involved in rMuV-induced innate immune response

To illustrate whether the altered m^6^A-landscape in both A549 andTHP1-iDCs is the host’s response to restrain the virus, or is MuV’s strategy to suppress the host’s innate immunity for more efficient viral reproduction, we first knockdown (KD) METTL3 by shRNA in A549 and THP-1, which was then used for THP1-iDCs induction. As the A549-shMETTL3 cells were infected by rMuV at the MOI of 0.2, less pIRF3, pIRF7, and RIG-I were induced at 16 and 24 hpi in shMETTL3 cells compared with the shN cells (Fig. 4a), meanwhile MuV viral protein N was mildly upregulated in either translation or transcription in A549-shMETTL3 (Fig. 4a, b). METTL3-knockdown also suppressed the transcription of type I and III IFNs and TNF-α, however, the IL-6 was significantly enhanced in both mock and infected cells (Fig.4c). In THP1-iDCs METTL3-knockdown suppressed the infection-induced phosphorylation of pIRF3 without affecting RIG-I, TLR7 and MuV N protein (Fig. 4d), and resulted in the impaired transcription of IFNs, IL-6, and TNF-α, as well as the expression of DCs maturation markers CD11c and CD86a, but partially rescued the HLA class II molecules, which was depressed by the infection (Fig.4f). It’s noticeable that METTL3 was also downregulated by the infection in the shN cells of both A549 and THP1-iDC (Fig.4a, d), like what happened in the corresponding wild type cells (Fig. 3a, b). To further confirm the m^6^A machinery effect on the MuV-induced iDC maturation, the FTO-targeting chemical inhibitor FB23-2^[35]^, were applied during THP1-iDC induction, and the cells were then infected by rMuV (Fig.4g). The FTO-inhibition was confirmed by the enhanced m^6^A abondance on the purified cellular mRNA (Fig.4h), and the FB23-2-treated iDC presented significantly enhanced CD11c, CD80, CD86 and HLA class I molecules, but depressed HLA class II molecules (Fig.4i). As the live rMuV-infection resulted in the decreased expression of multiple m^6^A machinery proteins (Fig. 3a, b, Fig.4a, b), which in turn may alter the virus reproduction (Fig.1d-f), to simply explore how the host’s m^6^A affects the rMuV-induced innate immune, MuV’s vRNA was purified and transfected into A549-siMETTL3 or A549-siN at a dose of 5 copies gRNA per cell, with the control of or poly(I:C) and PBS. It’s interesting to find that METTL3’s effect on innate immunity is totally different from the viral infection; the vRNA-induced innate immunity was further upregulated by METTL3-deficiency, except for IFITM123 (Fig.4j, k). In THP1-iDC, the impaired METTL3 also promoted the vRNA-initiated innate immunity in either IFNs and cytokines transcription, or the maturation markers expression (Fig.4l-n). Thus we demonstrated that in A549 and THP1-iDC the amount of host m^6^A deposit negatively modulate the type I and III IFN response against the exogenous viral RNA. However, as the live rMuV itself modulates the host m^6^A machinery, and may use viral proteins to affect multiple signaling pathways for the purpose of more robust reproduction or innate immunity escape, like many other RNA viruses do in their host cells^[36]^, so the observed m^6^A-modulated phenotypes in the context of infection and transfection are not consistent.

**Fig. 4.**
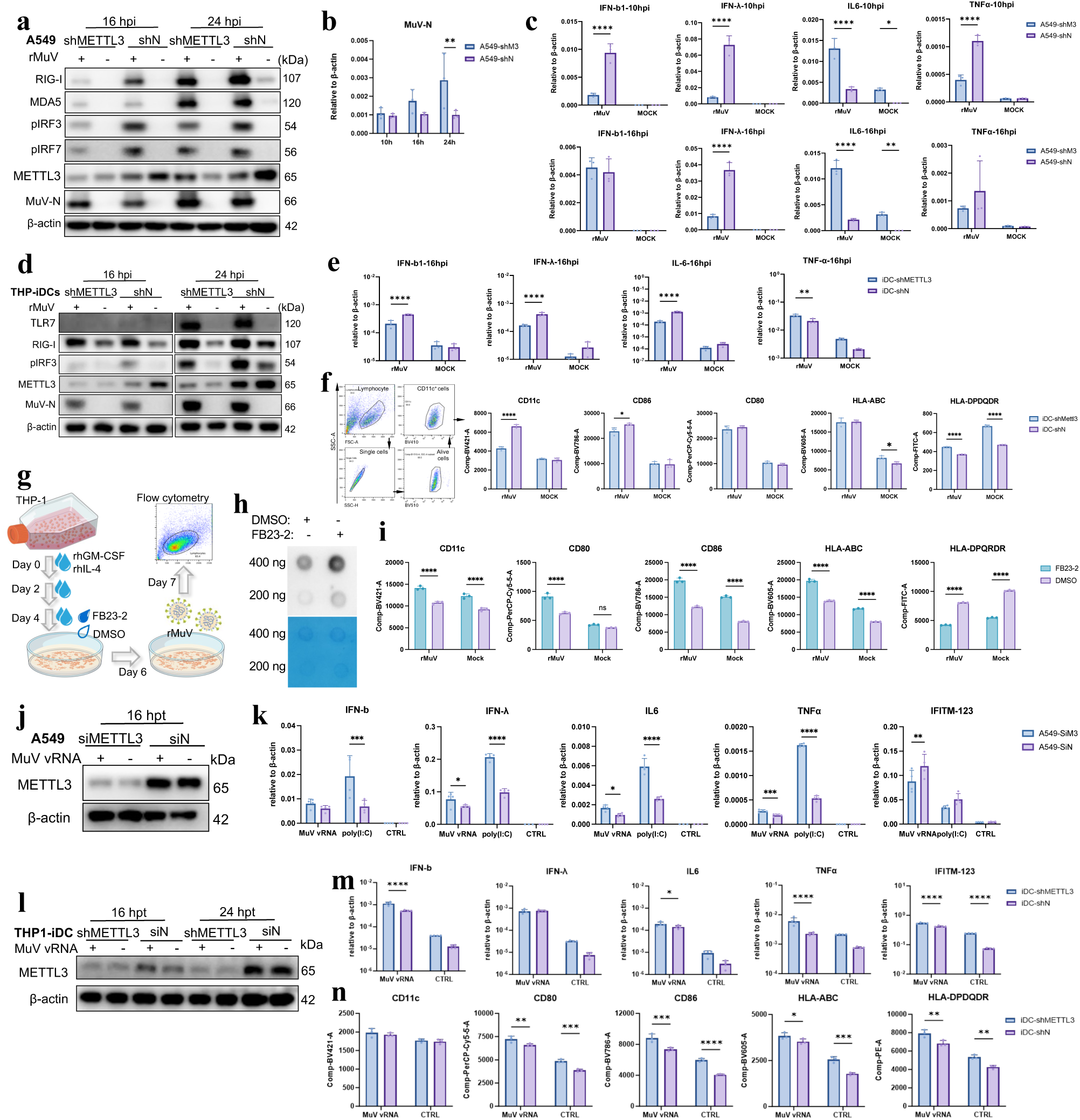
The host cells’ m^6^A-machinary is involved in rMuV-induced innate immunity. **a-c**. The A549-shMETTL3 cells were infected with rMuV at the MOI of 0.2, with the control of A549-shN. The cell lysate and total RNA were collected at the indicated time points for immunoblotting (**a**), viral N gene (**b**), IFNs and cytokines transcription quantification (**c**). **d-f.** The THP1-iDC-shMETTL3 cells were infected with rMuV at the MOI of 0.2, with the control of shN cells. The cell lysate and total RNA were collected at the indicated time points for immunoblotting (**d**), IFNs and cytokines transcription quantification (**e**), and the cell suspension was collected for maturation evaluation by flow cytometry. The cells were gated for lymphocyte-single cells-alive cells-CD11c-positive cells and then sub-gated for the indicated molecule-positive cells portions. The fluorescence strength for each single surface marker was generated by the software FlowJo (**f**). **g-i.** The chemical FB23-2 (10 μM) was added into the THP1-iDC induction culture medium at day 4 to specifically inhibits FTO, and the iDC was used for rMuV-infection at day 6 (**g**). The host cells’ mRNA was purified from total RNA and quantified, loaded onto the Hybond-N+ membrane and UV-linked for the following an antibody-based detection for m^6^A amount (**h**). The infected iDC was subjected to flow cytometry at 24 hpi for the maturation comparison between the control and FTO-inhibited cells (**i**). **j-k.** The A549 was transfected by siMETTL3 or siN 72 h prior to the transfection by rMuV vRNA (5 copies/cell). The cell lysate and total RNA were collected at the indicated time points for knockdown confirmation (**j**), IFNs and cytokines transcription determination (**k**). **l-n.** The THP1-iDC-shMETTL3 cells were transfected by rMuV vRNA (5 copies/cell), with the control of shN cells. The cell lysate and total RNA were collected for METTL3-deficiency confirmation by immunoblotting (**l**), IFNs and cytokines transcription quantification at 16 hpi (**m**), and the cell suspension was collected for maturation evaluation by flow cytometry at 24 hpi (**n**). The above data are from independent three experiments and the representative one is presented (**b, c, e, f, i, k, m, n;** mean ± S.D.). \**p* < 0.05, ** *p* < 0.01, *** *p* < 0.001, **** *p* < 0.0001 (Two-way ANOVA).

### m^6^A-deficient MuV promotes innate immune response by PRRs recognition

As the m^6^A-deficient (m^6^A^def^) virion RNA from multiple NNS viruses, including RSV, hMPV, and VSV, was proved to trigger more intense innate immunity through higher affinity to RIG-I^[11,12,21,22]^. We were wondering if MuV-triggered innate immunity is also modulated by the m^6^A scarcity on vRNA. To reduce m^6^A abundance in vRNA, STM2457-treated Vero cells were used to produce rMuV in bulk for vRNA purification (Fig.5a). Having been testified for m^6^A deficiency by dot blot (Fig.5b), the m^6^A^def^ vRNA was introduced into A549 cells by transfection and resulted in an elevated expression of RIG-I, MDA5 and pIRF3, as well as promoted transcription of IFN-β and IL-6, as compared to the vRNA from DMSO-treated cells (Fig.5c, d). Again we noticed that in THP1-iDC, the m^6^A^def^ vRNA induced promoted pIRF3, but no obvious enhancement in the expression of RIG-I and MDA5 (Fig.5e) line in A549 (Fig.5c), similar to these PRRs expression characteristic of not being affected by host m^6^A profile (Fig.4d), suggesting that in iDC the wild type vRNA-stimulated type I IFN is saturated for positive feedback on the PRRs, and a cell type-specific sensitivity of RLRs to IFN. The m^6^A^def^ vRNA also activated higher IFNs and pro-inflammatory cytokines in iDC, but downregulated TGF-β (Fig.5f), the differentiation factors of inhibitory Treg cells^[37]^, and promoted the maturation of iDC (Fig.5g), implying that the function of priming T cells might be more efficiently activated by the m^6^A^def^ vRNA. To illustrate the innate immunity signaling pathway activated by the m^6^A^def^ vRNA, the PRRs affinity assay was performed in HEK293T cells using RIG-I-, MDA5-, TLR3-, and TLR7-immunoprecipitation (IP) followed by RT-qPCR to quantify the PRRs-bound vRNA copies. As expected that RIG-I has the highest affinity to the m^6^A^def^ vRNA according to the relative binding (IP/ Input), and TLR7 has the poorest affinity with no difference between m^6^A^def^ and wild type vRNA (Fig.5h). Besides RIG-I, MDA5, TLR3 and TLR8 also prefer to bind m^6^A^def^ vRNA rather than the wild type one, though these PRRs don’t acquire 5’ppp on ligand RNA. So we next used Vero-E6-shMETTL3 and Vero-E6-shN to produce rMuV (rMuV(shM3) and rMuV(shN), respectively) to test the ability of activating innate immunity in A549. At the same MOI of 0.2 rMuV(shM3) initiated more robust type I and III IFN and cytokines, as well as ISG of IFITMs at 16 hpi, but the difference was eliminated gradually until 32 hip (Fig.5i). Collectively m^6^A-deficient MuV RNA is more efficiently sensed by PRRs of RLRs and TLRs and thus activates more intense innate immunity in both A549 and THP1-iDC, even more conveniently, the METTL3-defficient host cells-produced rMuV is also a stronger stimulus for production of IFNs and pro-inflammatory cytokines.

**Fig. 5.**
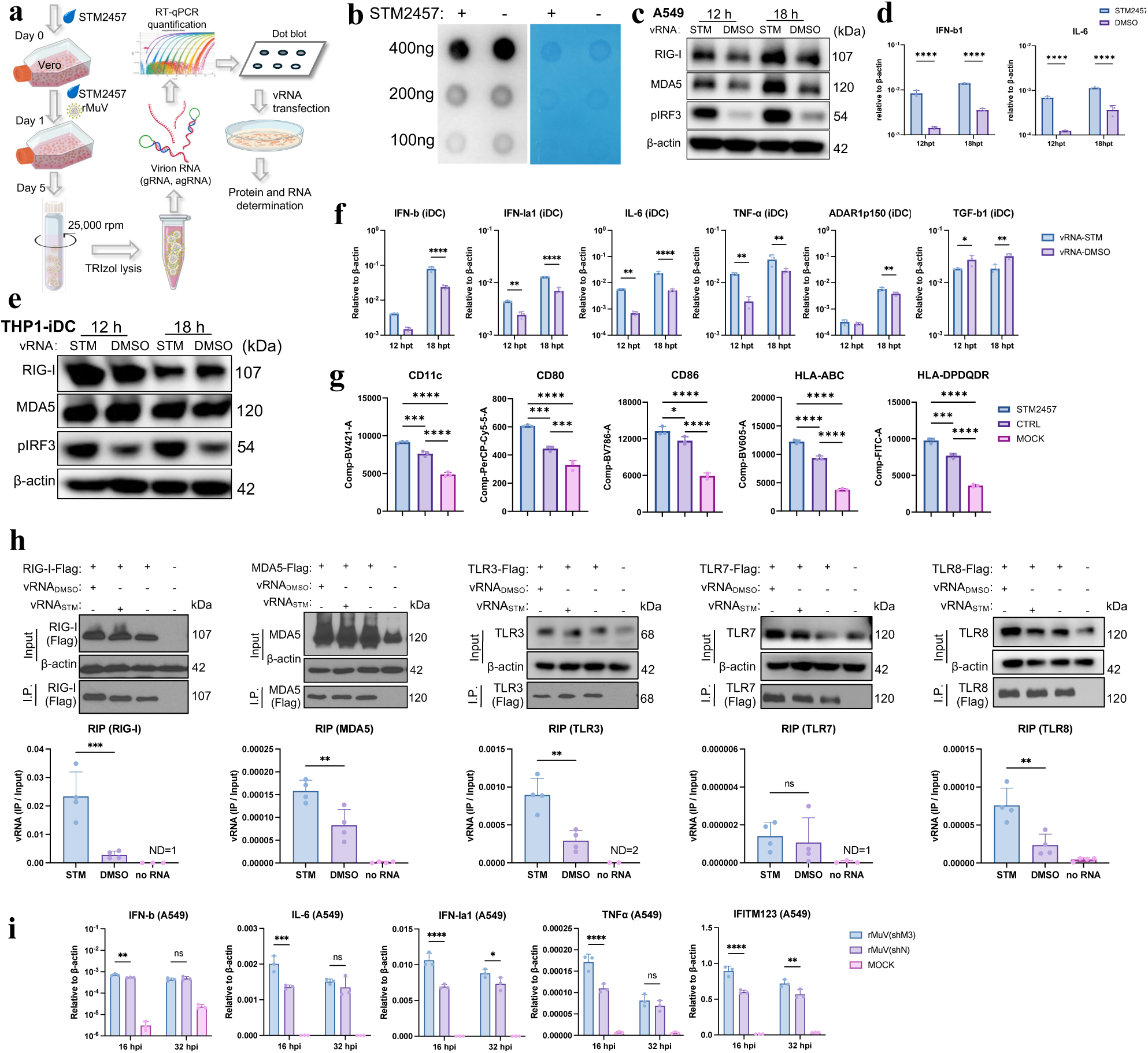
m6A-deficient MuV promotes innate immune response by higher affinity to PRRs. **a.** The chemical STM2457 was applied to inhibit METTL3 activity at 10 μM for 24 h incubation and then used as host cells to amplify rMuV in bulk for the following ultracentrifugation. The purified virion was used to extract vRNA, which was quantified for RNA copies and labeled as vRNA_STM_ and vRNA_DMSO_ for the subsequent performance. **b-d.** The vRNA_STM_ and vRNA_DMSO_ were subjected to m^6^A amount evaluation by dot blot (**b**). The vRNA-transfected A549 (5 copies/cell) was collected at 12 and 18 hpi for the innate immune response evaluation by immunoblotting (**c**) and RT-qPCR (**d**). **e-g.** The vRNA-transfected iDC (2.5 copies/cell) was collected at 12 and 18 hpi for cell lysate and total RNA for the innate immune response evaluation by immunoblotting (**e**) and RT-qPCR (**f**), and the cell suspension was collected for maturation evaluation by flow cytometry at 24 hpi (**g**). **h.** The HEK293T cells overexpressing Flag-tagged PRRs (RIG-I, MDA5, and TLR3/7/8) were transfected with the vRNA_STM_ or vRNA_DMSO_, at the dose of 100 RNA copies/cell. At 2 hpt the cells were lysed and subjected to Flag-tag-targeting immunoprecipitation (IP). The vRNA, which was pulled down by each PRR, was calculated for input and IP by RT-qPCR, and the binding efficiency (IP/Input) of each PRR molecule was calculated and compared between the DMSO- and STM2457-treatment. **i.** The Vero-E6-shMETTL3- or Vero-E6-shN-produced rMuV was used to infect A549 cells at the MOI of 0.2, and the transcription of IFNs, cytokines and ISG was determined by RT-qPCR at 16 and 32 hpi. The above data are from independent three experiments and the representative one is presented (**d, f, g, h, i;** mean ± S.D.). ns: *p* > 0.05, \**p* < 0.05, ** *p* < 0.01, *** *p* < 0.001, **** *p* < 0.0001 (Student’s T test, one-way ANOVA, and two-way ANOVA).

## Discussion

Multiple negative-strand RNA viruses (*e.g.* VSV, hMPV, hRSV, SeV and IAV), as well as positive-strand RNA viruses (*e.g.* SARS-CoV-2, ZIKA, DENV, PEDVnd HCV), have been discovered to carry m^6^A, which modulates either virus replication or host immune response through impairing the recognition by RIG-I^[31,38–42]^. As an ingredient of MMR vaccine, multiple strains of MuV have been used to inoculate young children for decades by injection method. Our previous work on COVID-19 vaccines inspire the application of MuV as an *in.*-dosed viral vector. But the innate immunity it induced in the epithelial or immune cells is not illustrated clearly, though the predominant involvement of RIG-I-MAVS axis is proved in mouse primary ovarian granulosa cells, sertoli cells and leydig cells^[43,44]^. Whether the RIG-I is also the predominant PRR in epithelial or immune cells, and whether the MuV-induced innate immunity is also deeply regulated by the viral RNA-carried m^6^A, or even the host m^6^A landscape, are not yet explored to date. We integrated a single-base resolution GLORI-seq to mapped the m⁶A landscape of a recombinant MuV JL2 strain. GLORI-seq is a newly emerged technique, which discovered numerous m^6^A peaks spatially enriched in the viral N, P, and HN regions (Fig.1, Table 3). Besides the specially uneven distribution, GLORI-seq-concluded motifs (MAACA and UAAAGW) don’t strictly conform to the canonical RRACH consensus, suggesting that viral RNA structure, replication intermediates, or polymerase-associated complexes may changes RNA accessibility and thus influence site selection, like the reported dynamic m^6^A modification in HIV-1, SARS-CoV-2, Japanese encephalitis virus (JEV), and Coxsackievirus B3 (CVB3), due to different mRNA structure and accessibility according to infection stages^[40,45–47]^. Functionally, depletion of METTL3 enhanced MuV replication in Vero-E6 cells, which lack a functional type I interferon system, demonstrating that m⁶A intrinsically suppresses viral propagation independent of antiviral signaling. Notably, RNase protection assays revealed that both gRNA and agRNA can be encapsidated, as what we’ve observed in human metapneumovirus (hMPV)^[11]^, and that nucleocapsid assembly efficiency is negatively regulated by m⁶A, which might modulate viral RNA secondary structure, affect polymerase processivity, or modulate interactions with viral/host RNA-binding proteins. This observation aligns with reports in several RNA viruses (e.g. PEDV, ZIKA, SARS-CoV-2, and HCV ), where m⁶A deposition restricts viral RNA stability or translation, or even the virus particle assembly^[40,42,48,49]^. Considering there’re much more high confident m⁶A peaks on MuV agRNA (Table 3) and the encapsidation of agRNA is nearly 0.5-fold of gRNA in the context of either normal or depleted METTL3, it implys that the m⁶A modification reduces the accessibility of agRNA to N protein, thus holding the plus strand RNA out from virus particles.

Some viruses even alter the subcellular location, expression, or degradation of m^6^A machinery molecules to facilitate its own replication through multiple mechanisms^[32,38,50–52]^, such as viral matrix protein (M) of bovine parainfluenza virus type 3 (BPIV3) binds to the methyltransferase domain of METTL3 in the nucleus and facilitates its translocation to the cytoplasm^[38]^. Pseudorabies virus (PRV) and herpes simplex virus type-1 (HSV-1), however, trigger phosphorylation of components of the METTL3/METTL14/WTAP complex by their own viral kinase protein US3 to inhibit the complex activity^[50]^. Our data discovered MuV infection profoundly altered host m^6^A machinery such as METTL3/14, FTO, and ALKBH5, and extensively redistributed m^6^A sites in the host transcriptome. The rMuV-infection increased the number of m^6^A-modified transcripts in both A549 and THP1-iDC cells, in contrast to the global m^6^A loss in some other RNA virus infections^[32]^. Due to the method application limit, non-coding RNA data is not obtained from GLORI-seq, but we noticed there’re much more lost-and gained-m^6^A sites in iDC mRNA (MeRIP-seq) than in A549 mRNA (GLORI) (Fig.3c), which might due to the cell type difference and the method accuracy/ resolution. Moreover, m^6^A fold changes positively correlated with transcript abundance, including multiple genes involved in innate immune pathways (*e.g.* PRRs, IFNs, ISGs, pro-inflammatory cytokines and DC maturation molecules), but not all immune-related transcripts exhibited parallel enrichment with m^6^A (e.g., TNF-α and HLA class I molecules), indicating selective targeting rather than global methylation enhancement. The asymmetric regulation of TLRs between epithelial and dendritic cells further underscores the integration of epitranscriptomic regulation with cell-specific immune programming (Fig.3h, *Supplementary Fig.1d*).

In the innate immune-competent epithelial A549 and monocytic THP1-iDC, MuV infection triggered dramatic innate immunity through multiple activated cytosolic PRRs-initiated signaling pathway. Among tested PRRs, RIG-I senses the MuV vRNA most efficiently in a m^6^A-dependent manner, which is also observed in MDA5-, TLR3 and TLR8-binding (Fig.5h). In THP1-iDCs, additional induction of TLR7/8 (Fig.2h) suggests a broader repertoire of RNA sensing in antigen-presenting cells, consistent with the enhanced m^6^A pattern (Fig.3h) and the reported cell type–specific PRRs predominance^[53]^. Despite robust innate activation, MuV infection led to downregulation of HLA class II molecules, indicating a skewed or incomplete DC maturation state and a potential impaired CD4^+^ T cell priming. But interestingly the depleted METTL3 partially restored the HLA class II molecules, and consistently when the demethylase FTO was inhibited by the chemical compound FB23-2, the molecules were further downregulated (Fig.4f, i). However, the METTL3 depletion and FB23-2 application resulted in an opposite phenotype pattern in regard to other maturation biomarkers and IFNs in both iDC and A549, as compared to HLA class II molecules, most of which in our iDC data are rarely m^6^A modified (*Supplementary* .1d), so this opposite effects might be due to an indirect modulation by other inhibitory mechanisms but not m^6^A. Also we noticed that METTL3 deficiency didn’t impair the upregulation of MDA5 and IL-6 in A549, and not the TLR7 and RIG-I in iDC (Fig.4a, c, d), though we suppose their upregulation is m^6^A-related (Fig.3h). This suggests a resource tilts of m^6^A modification in the context of innate immunity activation, in another word some transcripts own m^6^A-deposit privilege even under the METTL3 scarcity.

However, as the viral proteins were excluded, the vRNA-induced innate immunity was regulated by the METTL3 in a totally different way, in which METTL3 is negatively correlated with the type I/III IFN response and iDC maturation (Fig.4k, m, n). This negative modulation is also reported in severe COVID-19 patient samples, showing reduced METTL3 expression concurrent with high inflammatory gene expression in host cells^[17]^, and consistent with the fact that m^6^A negatively regulates IFN-β by YTHDF2-mediated faster turnover of IFN mRNAs^[54]^. Based on the conflict phenotypes between the viral-infection and vRNA-transfection contexts, we speculate that the infection resulted in a much more complicated context, including host m^6^A machinery retreat, an initiative hijack of m^6^A machinery by the virus to favor its own replication, direct interaction between viral and host proteins, and other epitranscriptomic modifications (*e.g.* m^1^A, m^5^C)-involved virus-host interaction. Furthermore in viral-infections more PRRs were involved and lead to a more comprehensive signaling network, such as NOD-like receptors (NLRs)^[55]^, Dendritic cell-specific intercellular adhesion molecules-3 grabbing non-integrin (DC-SIGN, a C-type lectin receptor on dendritic cells)^[56]^ (Fig.3g, *Supplementary* .1b), and Scavenger Receptor A1 (SR-A1)^[57]^, those sense virus-induced mitochondrial ROS and lysosomal damage, or bind high-mannose glycans on viral envelope proteins, but whether these pathway are modulated by m^6^A is still lacking of evidence.

TLR agonists have been used *in vitro* and *in vivo* as adjuvants to enhance Th1-type immune responses to DNA vaccines, subunit vaccines, and live-attenuated viral vaccines^[58–60]^. In our previous work, we tried to apply the m^6^A-deficient vRNA as a “carry-on” adjuvant for hMPV and RSV. To this end multiple m⁶A motifs were removed from the gRNA and agRNA using a time-consuming mutagenesis, and the resulting recombinant viruses activated more robust innate and adaptive immune response in animal model^[11,12]^. However, it’s hard to choose which and how many m⁶A sites to be mutated among hundreds of detected sites, and each mutant needs to be evaluated for growth efficiency, infectivity, virulence, innate and adaptive immunogenicity *in vitro* and *in vivo*. Here in this work, the METTL3-deficient Vero-E6-generated m^6^A^def^ MuV vRNA presented good affinity to multiple PRRs and induced stronger type I/III IFN and proinflammatory responses in either A549 or iDC, accompanied by higher maturation biomarkers and reduced TGF-β in iDC (Fig.5c-h). Meanwhile the m^6^A^def^ rMuV particles also induced mildly enhanced innate responses in A549 during early infection and gradually converged with the wild type one. Thus to produce a vaccine candidate virus in METTL3-deficient cell lines or accompanied by chemical compounds should be a safe, time-saving and high-yielding approach, without the genetic modification-associated biosafety concerns. Furthermore the m⁶A modification level on progeny viral RNA can be restored to normal, avoiding prolonged stimulation and chronic inflammation by genetically changed m^6^A^def^ vRNA. For this purpose, comprehensive and systemic in vivo studies are necessary in the future.

Collectively, our data uncovered that during MuV infection m^6^A operates as a bidirectional regulator on virus replication and host innate immunity. The work extends the methodology to optimize MuV’s immunogenicity in the context of being immunogen or viral vector, and also raises a potential mechanism that a proper or side effect of vaccination, such as prolonged inflammatory, delayed antibody emerge or even no-response, is modulated by the patient’s/ acceptor’s m^6^A machinery status, other than the vaccine’s immunogenicity. Future studies dissecting reader protein involvement, structural determinants of methylation-dependent PRR recognition, and in vivo relevance will comprehensively clarify the contribution of m^6^A to MuV pathogenesis and antiviral defense.

## Acknowledgements

The authors appreciate the members of our laboratory for helpful discussion, and acknowledge the support from the Scientific Research Center, Hangzhou Medical College.

## Author contributions

SW: data collection and analysis, writing-original draft preparation. YZ, XZ, TP, TF, QH: data collection and analysis. WW: writing-original draft preparation. YC: writing-reviewing and editing. ML: supervising this project, conceptualization and methodology, writing-reviewing and editing.

## Data Availability

All data generated or analyzed during this study are available from the corresponding author.

## Consent for publication

All authors contributed to the article and approved the final manuscript.

---

**Supplementary Fig. 1.**
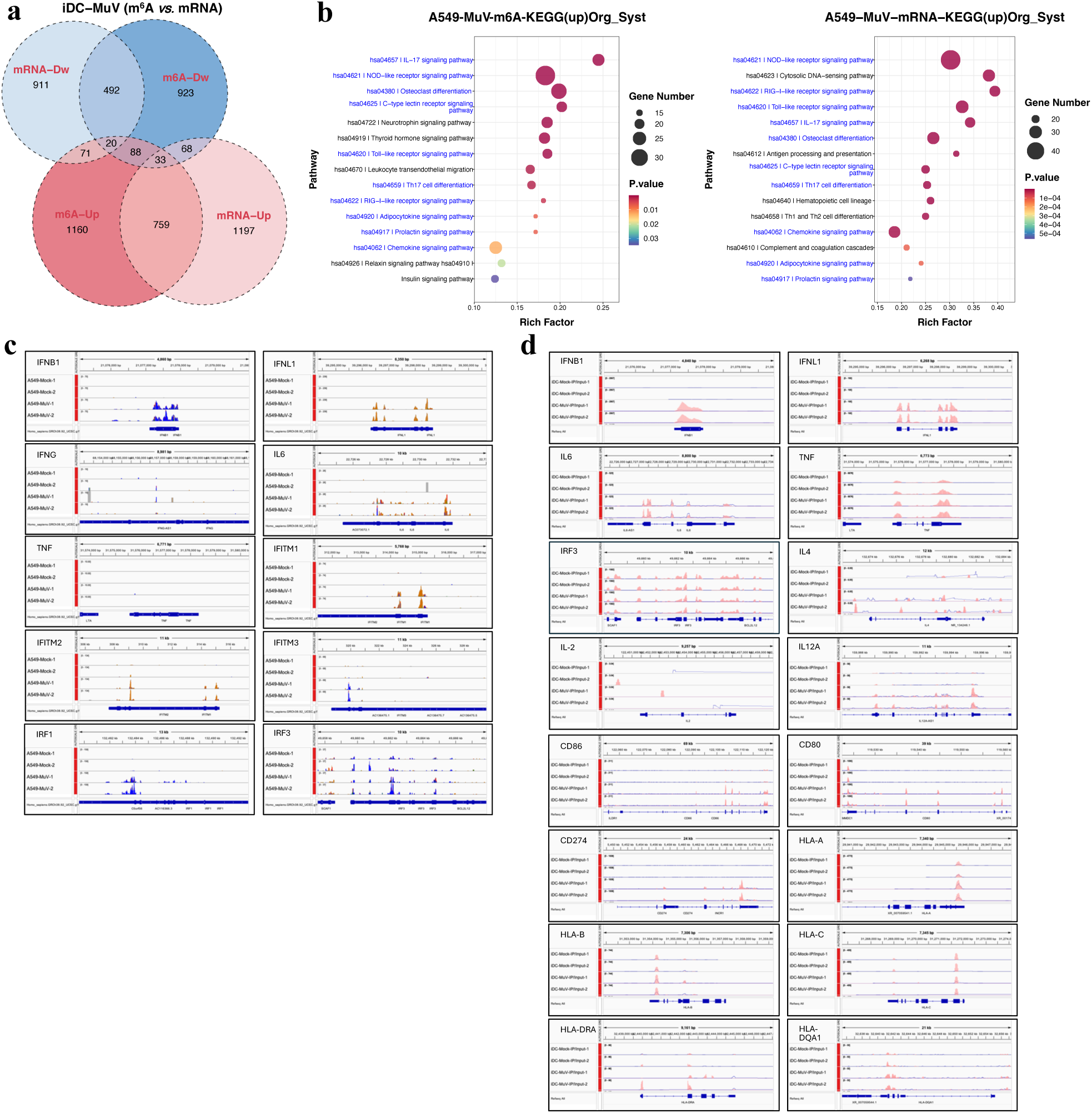
The rMuV induces parallel fold change of massive transcripts and the m^6^A-deposit. **a.** The fold change of mRNA and m^6^A in the rMuV-infected iDC is partially overlapped according to the gene ID and presented by the Venn plot, the area of each circle reflects the number of up- /down-regulated genes and m^6^A sites (threshold for significance: log2 fold-change > 1, and *p*< 0.05). **b**. In A549 the most enriched and upregulated KEGG pathways (top 15) in the category “Organismal Systems” are screened from the m^6^A-enriched (left panel) and the Input (right panel) mRNA according to the *p* values, the shared pathways in both datasets are labeled in blue **c-d.** The m^6^A enrichment in the indicated transcripts, which involve in innate immunity- (**c,** A549 cells) and DC maturation- (**d**, HTP1-iDC), were visualized in the software IGV, with a normalized confidence scale for both mock (two upper tracks) and rMuV-infected samples (two lower tracks).

